# AKT, NOTCH and GSK3β interact to trigger early myogenesis in vertebrate embryos

**DOI:** 10.1101/377804

**Authors:** Diane Lebrun, Pamela Rahal, Valérie Morin, Christophe Marcelle

**Affiliations:** Institut NeuroMyoGène (INMG), Université Claude Bernard Lyon1, Faculty of Medicine and Pharmacy, CNRS UMR 5310 - INSERM U1217, Lyon, France; Australian Regenerative Medicine Institute (ARMI), Monash University, Clayton, Victoria, Australia.

## Abstract

During early embryonic development, migrating neural crest cells expressing the NOTCH ligand Delta1 (DLL1) trigger the activation of NOTCH1 signaling in selected epithelial cells within newly formed somites. A key event in this process is a dramatic inhibition of GSK3β activity, initiated by the activation of NOTCH1 and that takes place independent of its transcriptional function. Here, we investigated the mechanism whereby NOTCH1 exerts its non-canonical function in somites. Using the activation of myogenesis as a read-out of the ability of NOTCH receptors to trigger transcription-independent responses in somites, we found that all NOTCH receptors (1-4) activate MYF5 expression and we showed that the RAM (RBPJ-Associated Molecule) domain of the NOTCH Intracellular Domain (NICD) is necessary and sufficient in this process. We then demonstrated that the NOTCH1 Intracellular Domain (NICD1) physically interacts in the cytosol with GSK3β and with the serine threonine protein kinase AKT. Activating AKT triggers myogenesis, likely via the inhibition of GSK3β. We found that AKT, in a dose-dependent manner, decreases the transcriptional activity of NOTCH, suggesting a role in the balance between the canonical and non-canonical functions of NOTCH. Altogether these data strongly support the hypothesis that transcription-independent function of NICD is a central mechanism driving myogenesis in early somites and suggests that, in this tissue, AKT, NOTCH and GSK3β interact in the cytoplasm to trigger a signaling cascade that leads to the formation of the early myotome in vertebrates.

## INTRODUCTION

During early embryogenesis, extensive tissue rearrangements and cell migration, associated with rapid cell fate changes, allow the formation of the tissues and organs of the future adult. A model where such complex issues are amenable to experimentation is the early formation of skeletal muscles in the chick embryo. Over many days of development, the medial border of the dermomyotome (DML) generates the first skeletal muscle cells that assemble into a primary myotome. This arises from a crucial cell fate decision: epithelial cells in the DML either self-renew or undergo myogenic differentiation, accompanied by an epithelial to mesenchymal transition (EMT) that allows their translocation into the primary myotome^1–4^.

We previously demonstrated that the activation of MYF5 and MYOD in the DML is dependent upon the transient activation of NOTCH signaling, itself triggered by a mosaic expression of Delta1-positive neural crest cells migrating from the dorsal neural tube, a mode of signaling we named “kiss-and-run” mechanism^4^. The molecular response that takes place within DML cells upon activation by neural crest cells was recently characterized^5^. Within DML cells, the “kiss-and-run” mechanism triggers a signaling module encompassing NOTCH, GSK-3β, SNAI1 and β-catenin and which couples the initiation of myogenesis in DML cells with their EMT (Figure 1A).

**Figure 1:**
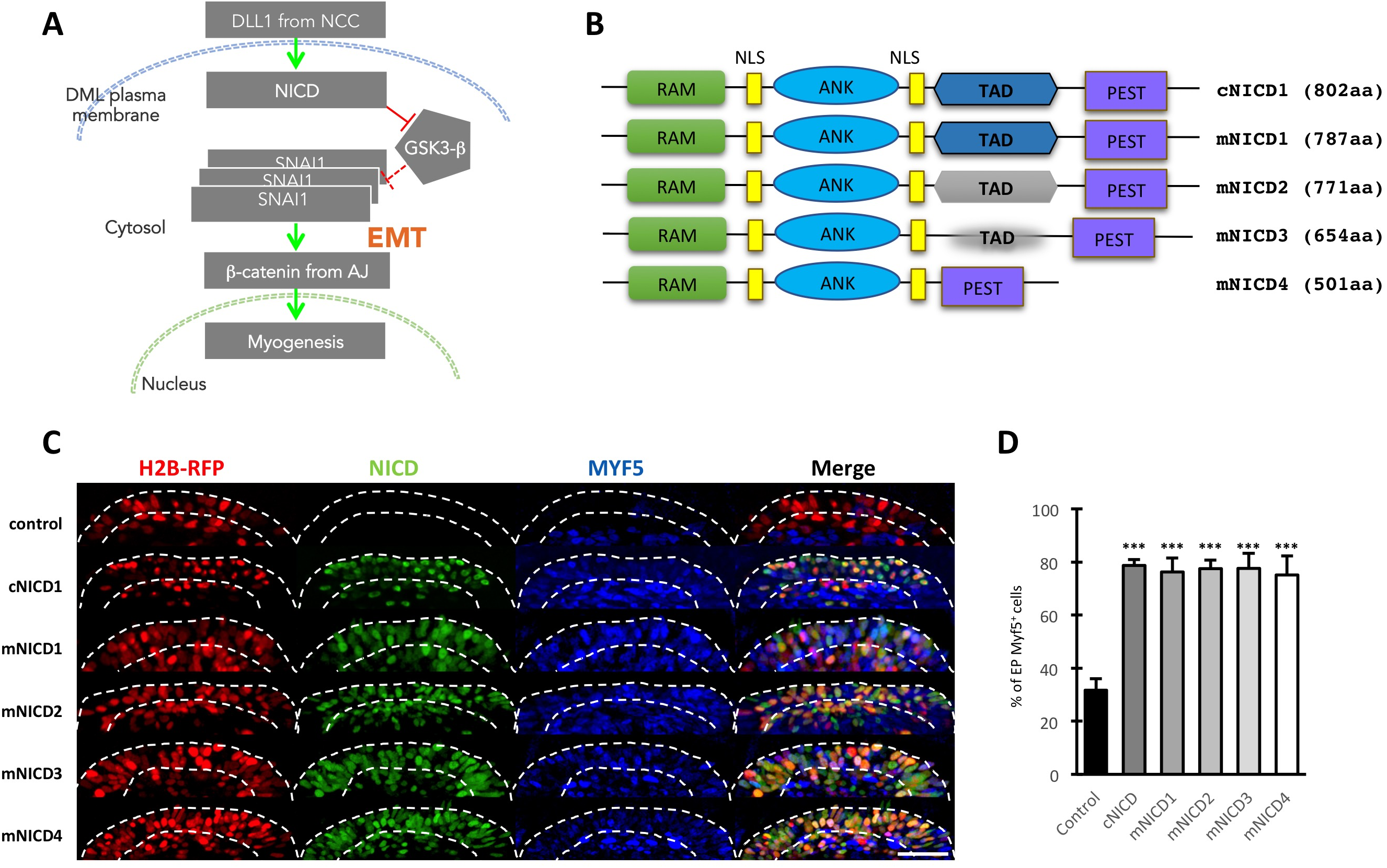
all NOTCH receptors activate MYF5 expression. (**A**) A simplified model representing the sequence of molecular events taking place at the DML during early myogenesis (adapted from ^5^). Delta1 (DLL1) from migrating neural crest cells (NC) activates NOTCH receptor in DML cells. The inhibition of GSK3β by NICD leads to the stabilization of Snail1 (SNAI1), which in turn triggers an EMT. As a consequence, βcatenin from the adherens junction (AJ) enters the nucleus where it activates myogenesis **(B)** Structure of the intracellular domains (NICDs) of the mouse NOTCH receptors (1-4) and of the chicken NICD1. All NICD constructs are tagged to allow detection (not shown). Indicated sizes are without the HA and Myc tags. **(C)** Confocal stacks of somites 6h after electroporation of the four mouse NICD isoforms, immunostained for c-myc (in green) and RFP (in red) to detect the electroporation marker (H2B-RFP). MYF5 expression is shown in blue. Dotted lines delineate the borders of the DML. (**D**) bar chart showing the percentage of electroporated MYF5^+^ cells compared to the total number of electroporated cells. ***:P<0,001 31,7% of cells electroporated with the electroporation marker express MYF5; 78,7% the cells electroporated with the chicken NICD express MYF5 and respectively 76,2%, 77,5%, 77,6% and 75,1% of the cells electroporated with the isoforms 1 to 4 of the mouse NICD express MYF5. Scale bar: 50*µ*m

In the vast majority of cellular processes, NOTCH utilizes a well-characterized “canonical” pathway initiated by the interaction of NOTCH with cell bound ligands (Delta-like or Jagged). This results in cleavage of NOTCH extracellularly by ADAM proteases, and intracellularly by the γ-secretase complex. This releases NICD, which translocates into the nucleus to interact with RBPJ, converting it into a potent transcriptional activator of downstream target genes^6, 7^. One of the remarkable features of the kiss-and-run signaling was that the activation of myogenesis by NOTCH seemed independent of its transcriptional function. This was based on the observation that a membrane-tethered NOTCH Intracellular Domain (NICD) displayed similar activity on myogenesis than NICD, while constitutively active and dominant negative forms of the NOTCH effector RPBJ had no effect on this process (Figure 1A). This led to the hypothesis that unconventional functions of NOTCH triggers early myogenesis^5^. Ligand-and transcription-independent functions of NOTCH have been reported in a growing number of cellular contexts and are collectively referred to as “non-canonical” NOTCH signaling^8–10^. NOTCH and WNT/β-catenin signaling are in fact known to interact through non-canonical mechanisms during development and in pathological processes, but in all cases studied, both pathways act antagonistically^11–15^. In contrast, we found that NOTCH promotes the β-catenin-dependent activation of myogenesis in early somites, suggesting that an atypical interaction is at work between these two major signaling pathways to initiate the myogenic program in developing somites.

To decipher the molecular events taking place downstream of NOTCH activation, we have now focused our attention on the mechanism inhibiting GSK-3β function. We demonstrated that *in vivo*, NICD, GSK3β and AKT work within a complex, in which AKT kinase activity is required to induce MYF5 expression, likely through its inhibitory role on GSK3β. Reinforcing this hypothesis, we found that SNAI1, a direct target of GSK3β, is a necessary step downstream of AKT in the chain of molecular events leading to myogenesis initiation. We present compelling evidence showing that NICD, GSK3β and AKT interact in the cytosol, suggesting a transcription-independent, non-canonical function of NOTCH in this cellular context. We observed that AKT negatively regulates NOTCH transcriptional activity, suggesting that the canonical and non-canonical NOTCH responses are mechanistically linked. Finally, we uncovered that all NOTCH receptors are able to activate MYF5 expression and we show that the RAM domain of NICD is necessary and sufficient in this process. Altogether these data shed a novel light on the molecular mechanisms whereby NOTCH regulates early myogenic differentiation in vertebrates and suggests that such signaling module could be active in a number of cellular contexts.

## RESULTS

### All four NOTCH receptors are capable of activating MYF5 expression

There are four NOTCH receptors in the mouse genome, while only three have been detected in the chicken genome^7, 16^. Since all receptors share similar canonical functions (e.g. complex formation with the transcription factor RBPJ), it is plausible that they could also share non-canonical, cytoplasmic activities when placed in similar cellular contexts. We speculated that the electroporation of early somites could serve as an experimental paradigm where this hypothesis could be tested. NOTCH1 was previously identified as the endogenous receptor responsible for triggering MYF5 (and MYOD) expression in the DML of chicken embryos through a transcription-independent mechanism^4^. We therefore used the induction of MYF5 expression as a read-out of the ability of other NOTCH receptors to display similar non-canonical functions. We electroporated the constitutively active forms of all four mouse NOTCH (mNICD1-4) in newly formed somites (Figure 1B). The chicken NICD1 (cNICD1) served as positive control while the electroporation vector alone (H2B-RFP) served as negative control (indicative of MYF5 expression in normal condition). Six hours later, we analyzed those embryos and we compared the expression of MYF5 in the DML of embryos electroporated with these constructs to that of wild type (WT) embryos. We observed that all four NICD induced a robust activation of MYF5 expression (very similar to that of cNICD1, Figures 1C-D), compared to controls. These data indicate that all NICDs share the same ability to activate myogenesis, likely through the signaling module that we previously uncovered, supporting the assumption that all NOTCH receptors are capable of displaying non-canonical function in specific cellular contexts.

### The RAM domain is necessary and sufficient for myogenic activation

We then determined which of the domains of NICD is responsible for its non-canonical function. NICD is composed of several domains, including the RAM (RBPJ Associated Module) domain, the 7 Tandem Ankyrin-like Repeats (ANK), the TAD (Trans-Activation Domain) and the PEST domain (Figure 2A^17^). The RAM domain and ANK repeats are thought to be necessary for NOTCH signaling through the canonical (RBPJ-dependent) pathway^18, 19^. The RAM domain presents high binding affinity for RBP-J and the ANK domain a low binding affinity for this molecule^20, 21^. Since both domains are implicated in protein-protein interactions, we wondered whether they could be involved in the molecular events leading to the induction of myogenesis in the cytosol of DML cells. Since the TAD domain is not present in mNICD4 (Figure 1A), while this molecule is capable of triggering myogenesis (see above), this excluded a role of TAD in early myogenesis. To test the function of the RAM and ANK domains, deletion constructs for each or both domains were generated and electroporated in DML cells of early chicken embryos (Figure 2A). Expression of MYF5 served as a read-out of their activity. As shown in Figures 2B and C, the deletion of the RAM domain significantly decreased the number of MYF5^+^ electroporated cells as compared to the WT NICD. In contrast, the deletion of the ANK repeats resulted in a percentage of MYF5^+^ electroporated cells similar to that obtained with the WT NICD. Finally, RAM and ANK deletion presented a significant decrease in MYF5^+^ cells, similar to that observed in the RAM-only mutant (Figures 2B-C).

**Figure 2:**
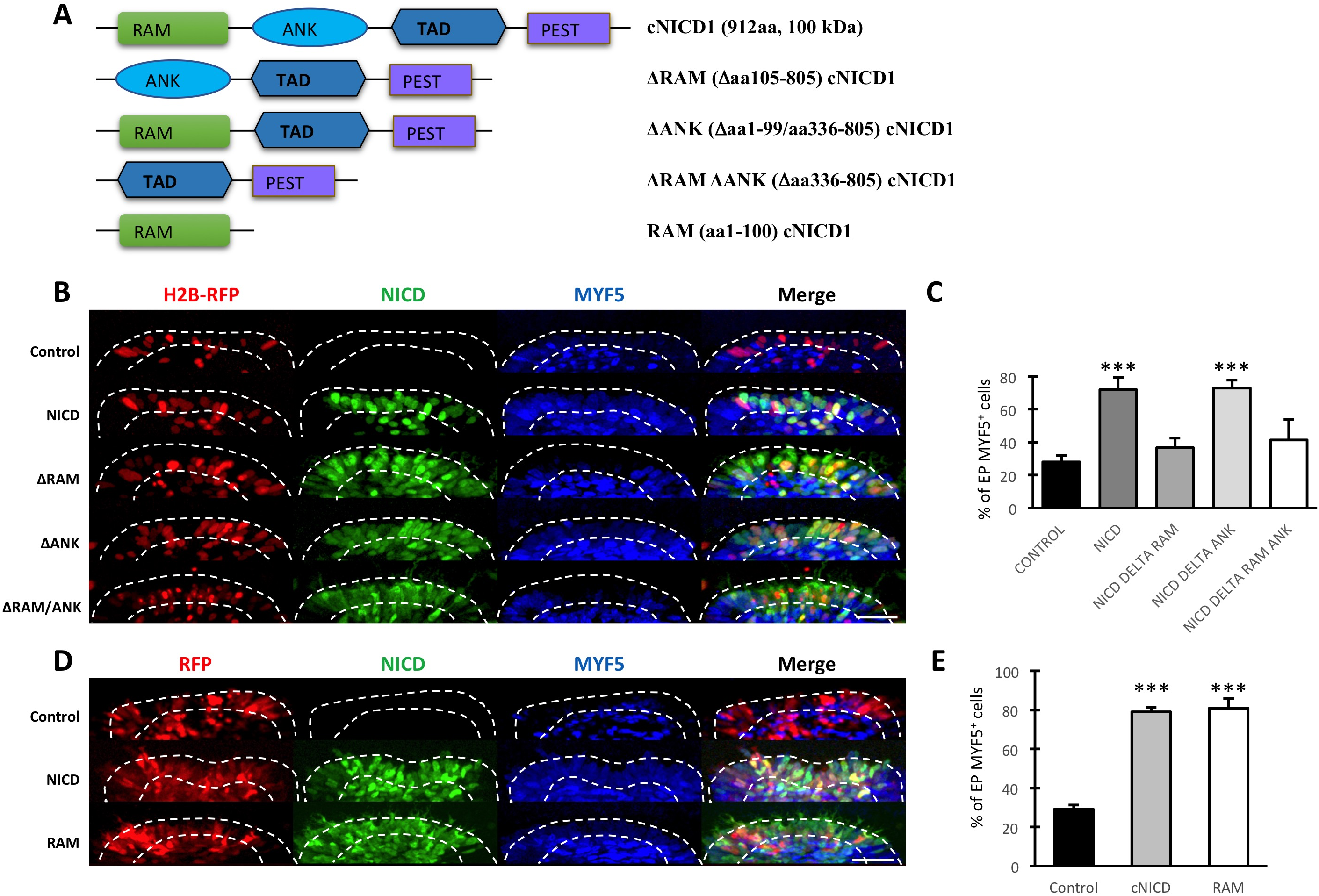
the NICD RAM domain is necessary and sufficient to induce Myf5 expression. (**A**) Schematics of the NICD mutants used in this experiments. All NICD constructs are tagged to allow detection (see Material and Methods). Size of cNICD1 (912 aa) takes the HA and Myc tags into account. Amino acid numbering is derived from the chicken NOTCH 1 (GenBank accession # XP_015134811.1). **(B)** Confocal stacks of somites 6h after electroporation of mutant forms of cNICD1 (in green) together with an electroporation marker (in red). MYF5 expression is shown in blue. **(C)** Bar charts showing the percentage of electroporated MYF5^+^ cells. 28,0% of electroporated cells express MYF5 in controls, 71,9% with NICD and 36,7%, 71,9, and 41,4% with a NICD missing the RAM domain, the ANK domain, and both the RAM and ANK domains, respectively. **(D)** Confocal stacks of somites after electroporation of either cNICD1 or the RAM domain of cNICD1 (in green), together with an electroporation marker (in red). MYF5 expression is shown in blue. **(E)** Bar charts showing the percentage of electroporated MYF5^+^ cells. 29,0% of electroporated cells express MYF5 in controls, 79,0% with chicken NICD and 80,8% with the RAM domain only. ***:p<0,001. Scale bars: 50*µ*m. Dotted white lines indicate the borders of the dorsomedial lip of the somite. Scale bars: 50*µ*m

Since we had previously shown that NICD or a membrane tethered NICD (CD4-NICD) comparably activate MYF5 expression in DML cells^5^, we tested whether the same would apply to the NICD deletion mutants. We have shown previously that the fusion of CD4 to NICD results in a molecule that does not enter the nucleus and is unable to activate NOTCH reporters^5^. We found that the deletion of the RAM domain from the CD4-NICD abrogated the induction of MYF5 expression observed after electroporation of the CD4-NICD alone. On the contrary, the deletion of the ANK repeats had no effect (Figure 2 - figure supplement 1 A-C). Together these results suggest that the RAM domain of NICD is necessary for activation of myogenesis in the cytosol of DML cells of the chicken embryo.

We next determined whether the RAM domain is sufficient to induce MYF5 expression in DML cells. To demonstrate this, we constructed a plasmid that encodes only the RAM domain (Figure 2A) and we electroporated it in DML cells. We observed that, compared to non-electroporated control embryos, the RAM domain was sufficient to increase the percentage of MYF5^+^ cells to a similar level to that observed after electroporation of the WT NICD (Figures 2D-E).

Altogether, these results suggest that the RAM domain of NICD is necessary and sufficient to trigger the chain of molecular events leading to the induction of myogenesis in the chicken embryo. Since we have shown previously that RBPJ is not involved in myogenesis in the DML^5^, this suggests that the RAM domain of NICD likely interacts with other protein(s) to mediate its pro-myogenic activity.

### NICD physically interacts with GSK3β and AKT in the cytoplasm of muscle progenitors

Using a reporter for GSK-3β activity, we have previously shown that both NICD and a membrane-tethered CD4-NICD induce a dramatic inhibition of GSK-3β activity^5^. Although previous studies suggested that GSK-3β binds to NOTCH to activate or inhibit its transcriptional activity^22, 23^, the mechanism whereby NOTCH could regulate GSK-3β activity is unknown. As a first step towards this aim, we tested whether NICD and GSK-3β physically interact *in vivo*. We performed co-immunoprecipation (co-IP) experiments on total protein extracts of somites electroporated with tagged (myc) NICD and CD4-NICD. We analyzed the embryos six hours after electroporation. This focuses on primary events resulting from expression of the constructs. About 40-50 electroporated embryos per condition were tested and the experiments were repeated 3 times. Protein extracts (1mg/lane) showed equivalent amounts of GAPDH protein in all samples (Figure 3A). Similar amounts of endogenous GSK-3β protein were also present in all protein extracts and the electroporated constructs (NICD and CD4-NICD) were also detected. Analysis of the immunoprecipitates showed that NICD and CD4-NICD co-precipitated GSK3β (Figure 3B). Quantification of three independent co-IPs suggests that NICD physically interacts with GSK3β in muscle progenitors present within the DML (Figure 3C). The observation that the NICD and the membrane-tethered NICD (CD4-NICD) co-precipitated GSK-3β, suggests that the physical interaction between NICD and GSK3β is taking place in the cytosol.

**Figure 3:**
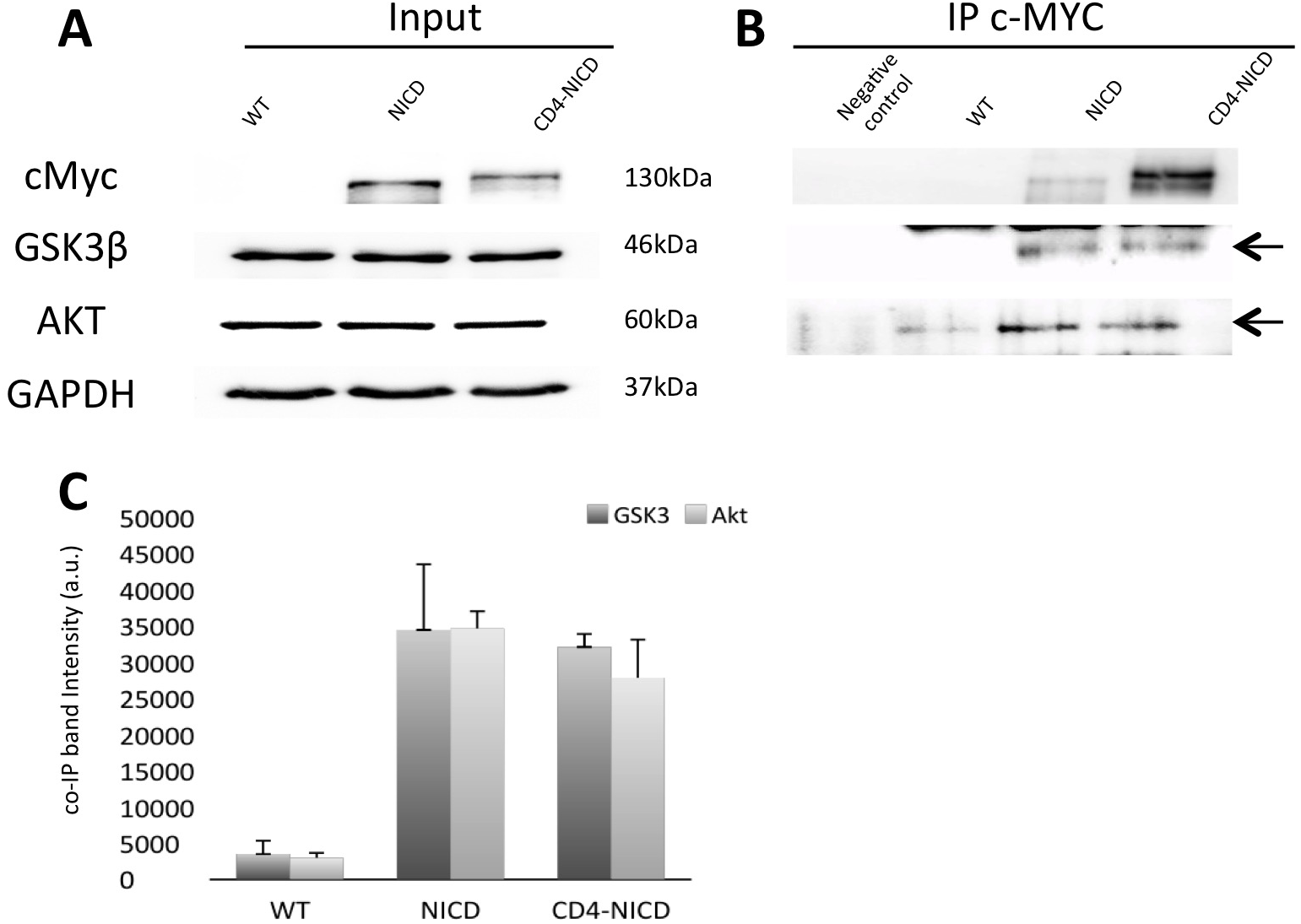
NICD physically interacts with GSK3β and AKT in the cytoplasm of muscle progenitors. Co-IP of endogenous GSK3β and AKT by exogenously expressed, tagged NICD and membrane tethered (CD4-)NICD. (**A**) Detection of GSK3β and AKT in total protein extracts (1mg) of DML electroporated with controls (WT), NICD and CD4-NICD plasmids. (**B**) 30 *µ*g of the protein extract were used for coIP with c-Myc antibody and reacted with GSK3β and AKT antibodies (for AKT, we used an antibody that recognizes all 3 AKTs). Negative control lane is a test of the co-IP beads specificity: NICD electroporated extract, no c-Myc antibody, and reacted with Magnetic coIP beads. (**C**) Bar charts representing the results (mesured in bands intensities) of 3 coIP experiments (as shown in B). GSK3β (dark grey) and AKT (light grey).

Since NICD does not display any kinase activity to inhibit GSK-3β, it is unlikely that NICD directly regulates GSK-3β activity. Therefore, it is probable that another kinase(s) is responsible for this effect. A well-documented GSK-3β modulator is the serine threonine kinase AKT (also known as Protein Kinase B)^24–26^, which phosphorylates and inhibits GSK-3β, notably in response to growth factor stimulation or to G-protein-coupled receptor (GPCR) activation.

We first determined whether AKT is co-expressed with NOTCH when myogenesis is first initiated in developing embryos, i.e. in the medial border of the dermomyotome of newly formed somites (DML)^4^. As in mammals, 3 AKT homologs (1-3) are present in the chicken genome. We performed in situ hybridization with probes for AKT1-3 on developing embryos and we observed that AKT1 is expressed in the DML at the time that muscle progenitors are generated in this structure, supporting the possibility that this molecule could play a role in this process (Figure 3 - figure supplement 1 A-B). This observation is coherent with data in mice, where AKT1 was found to be the main AKT expressed during embryonic development^27^.

In the somite extracts electroporated with the two NICD variants described above, we observed that endogenous AKT co-precipitated with NICD and CD4-NICD, suggesting that NICD and AKT proteins physically interact within the cytosol of DML cells (Figures 3A-C).

Together these results show that NICD directly interacts with GSK3β and AKT at the initial step of the signaling cascade leading to myogenesis. Furthermore, given the reported inhibitory role of AKT on GSK3β, our results suggest that AKT mediates the GSK3β inhibition needed to trigger the signaling cascade for induction of myogenesis in the DML.

### AKT induces MYF5 expression in a kinase-dependent manner

If AKT were to inhibit GSK3β in the DML, modulating its activity should affect myogenesis. To test this we electroporated the DML of developing chicken embryos with a construct coding for a wild type or a constitutively active form of AKT (CA AKT). Quantification of the proportion of electroporated cells expressing MYF5 showed that both WT and CA AKT expression led to a robust increase in the proportion of MYF5-positive cells (Figures 4A-B). This increase was alike the effect obtained after NICD over-expression in DML cells (see above and^4^). To verify that the kinase domain is required for this effect we electroporated a kinase-dead form of AKT. In this case, the number of MYF5-positive cells was comparable to control embryos. Altogether, these data suggest that the kinase activity of AKT controls the activation of MYF5 expression in the DML.

**Figure 4:**
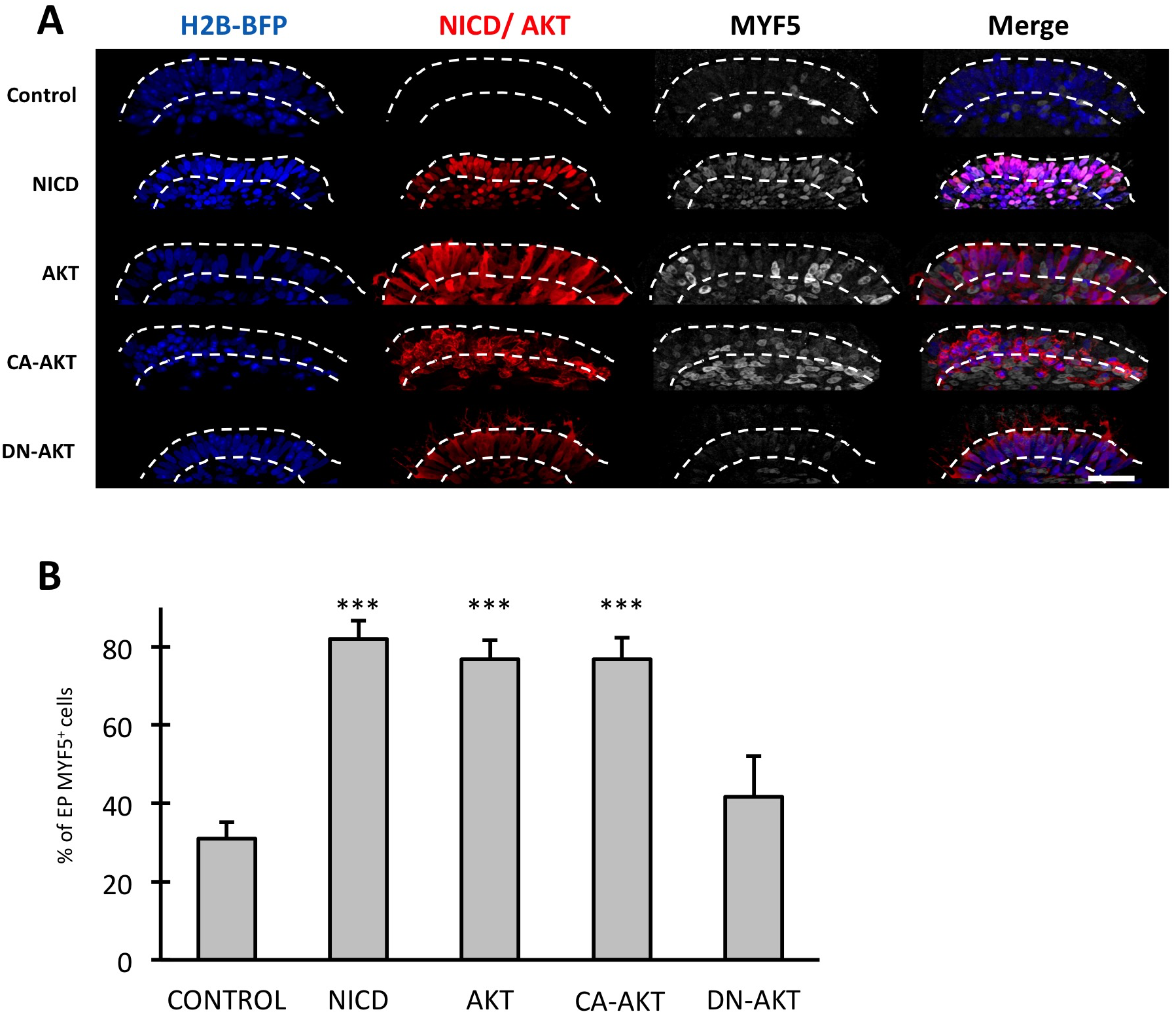
Akt induces myogenesis in a kinase-dependent manner. (**A**) Confocal stacks of somites 6h after electroporation of NICD or several AKT variants (in red) together with a nuclear electroporation marker (in blue). MYF5 expression is shown in grey. (**B**) Bar chart showing the percentage of electroporated MYF5^+^ cells in the DML. Controls electroporated with the marker (31,0% of electroporated cells express MYF5); 81,9% MYF5^+^ with NICD and 76,8%, 76,9, and 41,7% were MYF5^+^ after electroporation of ATK, Myr-AKT (CA) and DN-AKT. ***: p<0,001. Dotted white lines indicate the borders of the DML. Scale bar: 50*µ*m

### AKT regulates myogenesis through SNAI1

The result above is coherent with the hypothesis that AKT acts on GSK3β. However, AKT is a pleiotropic kinase with a multitude of known substrates. Therefore, it is possible that the activation of MYF5 expression we observed after activation of AKT was only partly due to its activity on GSK3β. To address this possibility, we tested whether blocking a downstream target of GSK3β would counteract the AKT-mediated activation of MYF5. A direct consequence of GSK3β inhibition in muscle progenitors is a stabilization of the transcription factor Snail1 (SNAI1)^5^. SNAI1 is a direct target of GSK3β^28^ and SNAI1 stabilization is a necessary step for activation of MYF5 in the DML^5^. We have co-electroporated AKT, together with a dominant-negative form of SNAI1 (DN SNAI1^5, 29^). We observed that the increase in MYF5 expression observed with AKT alone was abrogated when DN SNAI1 was co-expressed with AKT (Figures 5A-B). This confirms SNAI1 activation is a necessary step downstream of AKT-mediated MYF5 expression, and altogether reinforces the hypothesis that, within the time frame of these experiments, AKT regulates GSK3β activity in the DML.

**Figure 5:**
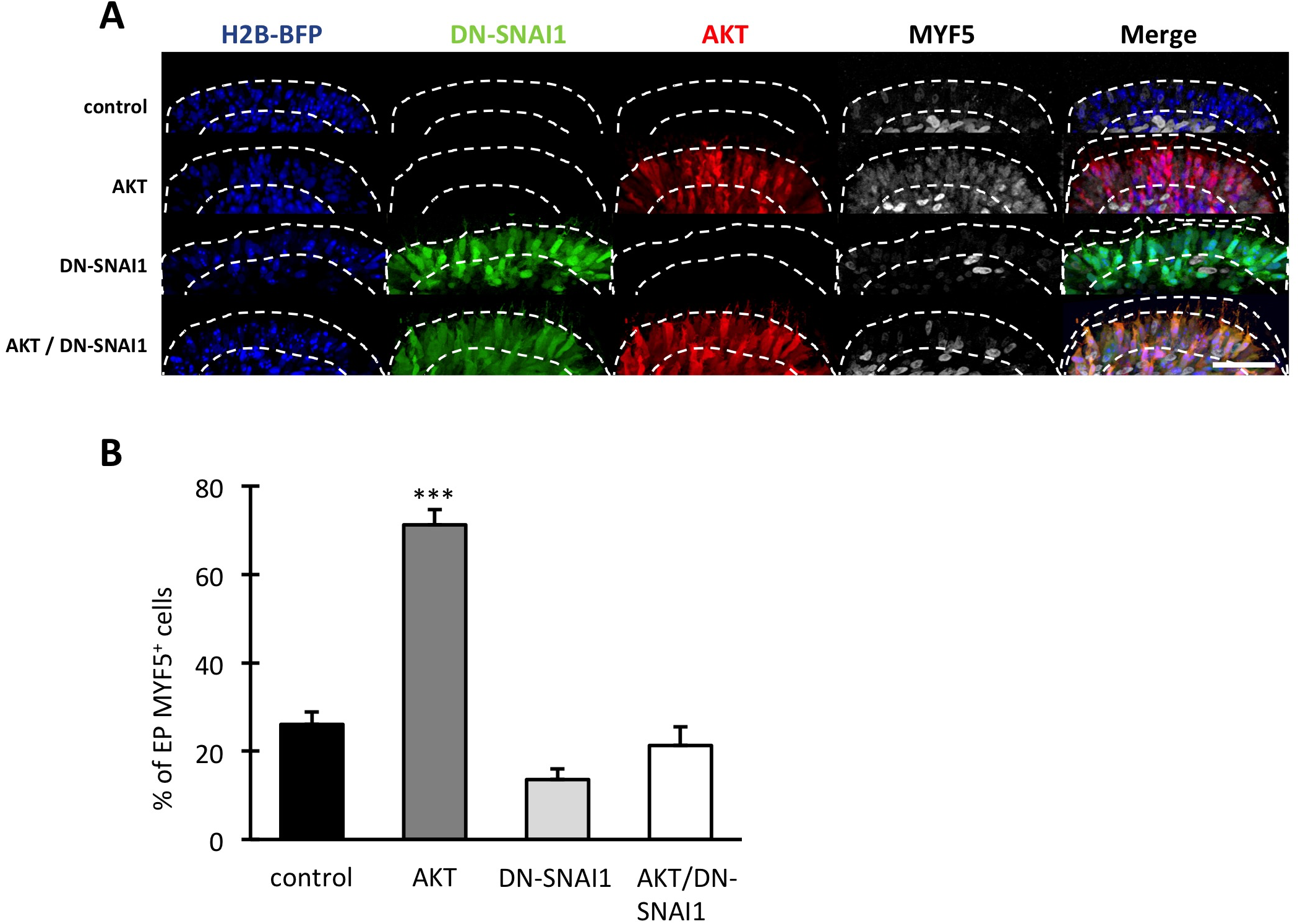
AKT regulates myogenesis through SNAI1 stabilization. (**A**) Confocal stacks of somites 6h after electroporation of AKT (in red), a dominant negative form of SNAI1^29^ (in green) or both plasmids, together with an electroporation marker (blue). MYF5 is shown in grey. Dotted white lines indicate the borders of the DML. Scale bars: 50*µ*m. (**B**) Bar chart showing the % of electroporated MYF5^+^ cells. Controls: 26,0% MYF5^+^; AKT 71,2% MYF5^+^; DN-SNAI1, 13,6% MYF5^+^; AKT & DN-SNAI1, 21,2% MYF5^+^. ***: p<0,001. Scale bar: 50*µ*m

### AKT, independent of its kinase activity, negatively regulates NOTCH transcriptional function

The data above strongly suggest that AKT is a novel component of the signaling module, upstream of SNAI1, which translates the activation of NOTCH by migrating DLL1-positive neural crest cells into an initiation of the myogenic program in DML cells. To test whether AKT may also regulate the canonical function of NOTCH, we co-electroporated increasing amounts of a tagged form of AKT together with a NOTCH reporter^4^. We observed that the activity of the reporter decreased with the increasing quantity of AKT (Figures 6A-B). The decrease in NOTCH transcriptional activity is likely independent of AKT kinase activity, since the co-electroporation of a dominant-negative (DN, i.e. kinase-dead) and a constitutively active mutant forms of AKT led to a similar decrease in the NOTCH reporter activity (Figures 6C-D). Finally, we found that the NOTCH transcriptional activity was similarly decreased when membrane-tethered forms of AKT and DN AKT were electroporated in DML cells (Figures 6E-F). Altogether, these data suggest that the balance between the transcriptional and non-transcriptional activities of NOTCH is mechanistically linked in DML cells. The mechanism whereby this regulation may take place is yet to be uncovered, but our data suggest that it may be due to an interaction between NICD and AKT that takes place in the cytosol.

**Figure 6:**
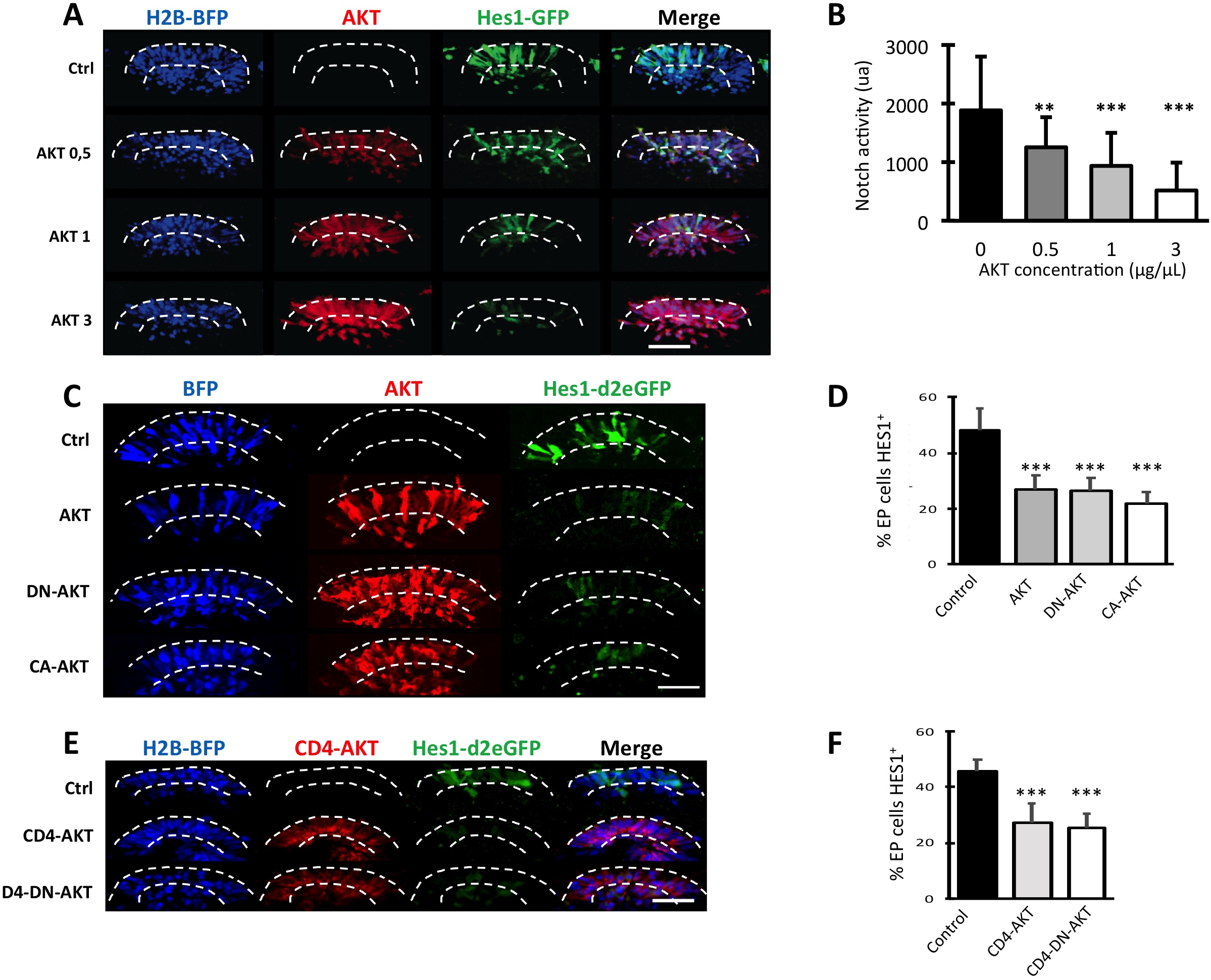
AKT negatively regulates NOTCH transcriptional function, independently of its kinase activity. (**A, C, E**) Confocal stacks of somites 6h after electroporation (in red) of AKT (A,C); a constitutively active form of AKT (CA-AKT) and a dominant negative form of AKT (DN-AKT, C); a membrane-tethered form of AKT (CD4-AKT) or of DN-AKT (CD4-DN-AKT, E). They were co-electroporated with a reporter of NOTCH transcriptional activity (Hes1-GFP^4^, in green) and an electroporation marker (in blue). Dotted lines indicate the borders of the DML. Scale bars: 50*µ*m. (**A**) Increasing amounts of AKT. (**C**) Electroporation of AKT, DN-AKT, CA-AKT. (**E**) Electroporation of CD4-AKT or CD4-DN-AKT. (**B**) Bar charts showing the decreasing transcriptional activity of NOTCH with increasing amounts of exogenous AKT. Relative intensities (in U.A.): 1884 in controls (no AKT); 1254 with 0,5*µ*g/*µ*L AKT; 937 with 1*µ*g/*µ*L; 517 with 3*µ*g/*µ*L. (**D**) Bar charts showing that 27,1% of DML cells electroporated with AKT express the NOTCH reporter; 26,4% with DN-AKT; 22,0% with CA-AKT; in controls, 48,1% of electroporated cells express the NOTCH reporter. (**F**) Bar charts showing 27,5% NOTCH reporter^+^ cells with CD4-AKT; 25,2% NOTCH reporter^+^ cells with CD4-DN-AKT; and 45,7% NOTCH reporter^+^ cells in controls. ***:p<0,001; **: p<0,005. Scale bars: 50*µ*m

## DISCUSSION

The purpose of this study was to characterize the mechanism whereby NOTCH plays non-canonical (ligand-dependent, but transcription-independent) functions in the DML. The demonstration that NOTCH, AKT and GSK3β physically interact in the cytosol of DML cells provides direct evidence for a role of NICD in the cytoplasm of those cells (Figure 7). It also identifies the binding of GSK3β to NOTCH as the initiating event, immediately downstream of NOTCH activation by incoming neural crest cells, in the signaling module leading to the initiation of MYF5 expression in this tissue.

**Figure 7:**
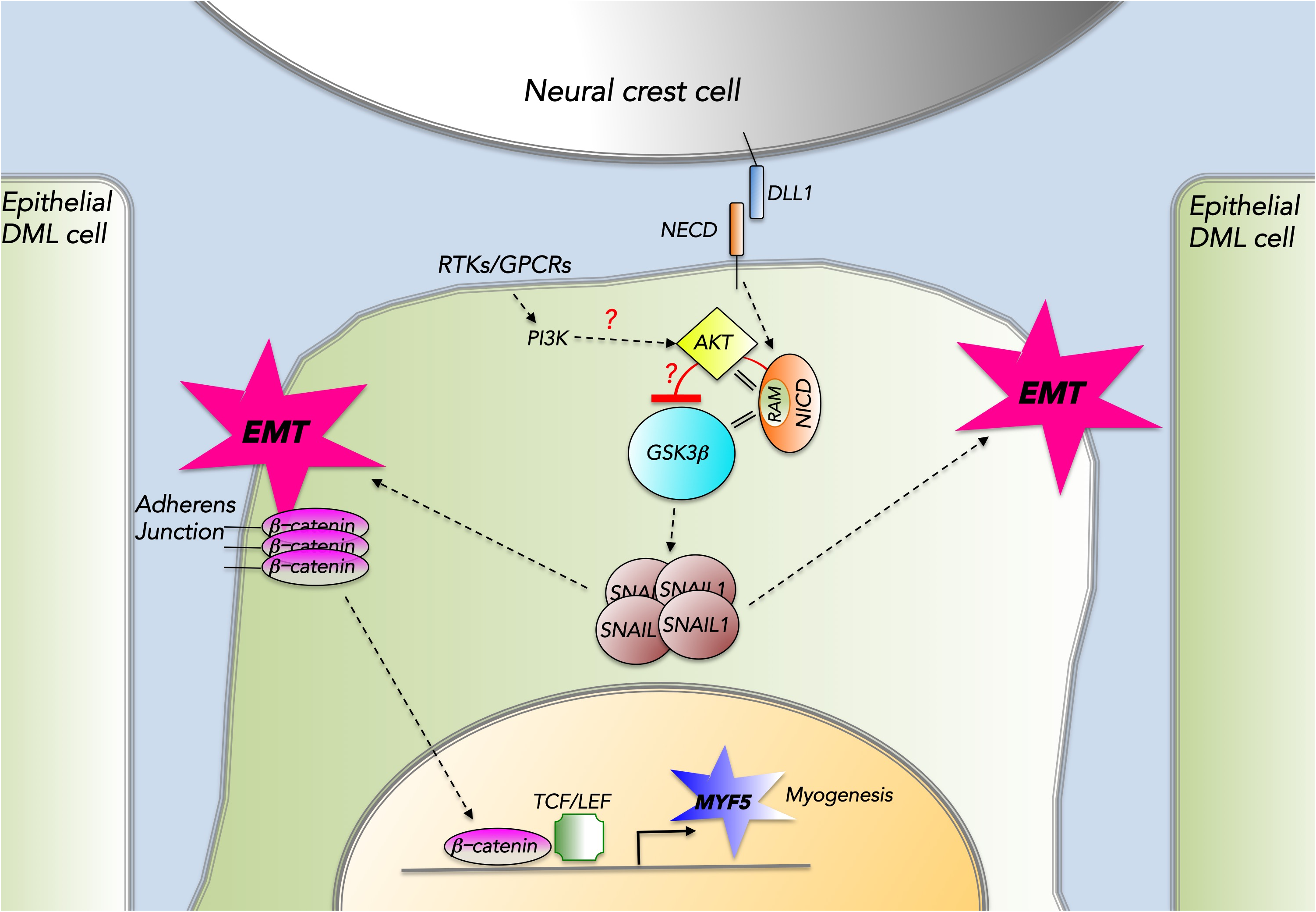
A model describing the main findings of this study. Contact between Delta1-positive neural crest and epithelial DML cells trigger the activation of NOTCH receptor in the latter. In the cytosol, this leads to the physical binding of NICD, through its RAM domain, to the serine-threonine kinases AKT and GSK3-β. The result of this interaction is a robust inhibition of GSK3-β, which leads to the stabilization of the EMT master gene SNAI1. AKT is an integral part of the signaling module leading to myogenesis, since it’s activation triggers MYF5 expression in a SNAI1-dependent fashion.

The identification of AKT in a protein complex with NOTCH suggests a possible mechanism whereby NOTCH could inhibit GSK3β activity: GSK3β being a major target of AKT^24–26^, it is tempting to postulate that its activity is inhibited by AKT within a protein complex encompassing all three molecules. AKT is a pleiotropic kinase that acts on a number or targets in various cellular contexts. In muscles, AKT is a major regulator of muscle mass through the mTOR and FoxO anabolic and catabolic pathways^30^. However, the observation that SNAI1 activation is a necessary step downstream of GSK3β and AKT supports a model whereby, within the short time frame of our experiments in DML cells, AKT regulates myogenesis solely through its action on GSK3β (Figure 7).

Interestingly, it was shown that the phosphorylation of NOTCH by GSK3β positively regulates canonical (transcriptional) NOTCH signaling *in vitro*^23^. It is thus possible that this explains the decrease in NOTCH transcriptional activity we observed with increasing AKT concentration (Figure 6). AKT and GSK3β would then act as molecular switches between canonical and non-canonical NOTCH signaling.

AKT is itself activated by the stimulation of receptor tyrosine kinases (RTK) or G-protein-coupled receptors (GPCR). Binding to their cognate ligand leads to plasma membrane recruitment and activation of the phosphoinositide 3-kinase (PI3K) that in turn activates AKT^31^. The uncovering that AKT plays a role in the signaling module triggering myogenesis therefore suggests that additional inputs from RTKs and/or GPCRs into the pathway leading to the activation of the myogenic program may play a role during this process. The nature of this signal(s) is only speculative at this point. Many FGFs (which signal through FGFR1-4 RTKs) are robustly expressed by the growing myotome, directly adjacent to the DML^29^. Frizzled (GPCR) that transduce WNT signals carried by migrating neural crest cells or secreted by the overlaying ectoderm are also present in the DML^3, 5, 32, 33^. It was also suggested that Sonic Hedgehog (SHH, whose role in early myogenesis has been debated^34^) activates AKT phosphorylation^35, 36^. Whether in the DML SHH, FGFs, WNTs or other signals synergize with NOTCH signaling *via* AKT to trigger early myogenic differentiation remains to be shown (Figure 7).

Finally, the observation that the NICDs of all four NOTCH receptors similarly activate myogenesis in the DML strongly suggests that they do so through similar, non-canonical mechanisms. The widespread expression of AKT, GSK3β and the NOTCH receptors suggests that similar interactions between those molecules could take place in many cellular contexts where cell fate changes are associated with an EMT.

Since NOTCH is such an extensively studied molecule, it may seem surprising that such interaction of NOTCH with widely expressed kinases was never reported. It is possible that the experimental paradigm we are using (analyses of tissues 2-3 hours after constructs initiate expression) allows detecting events immediately downstream of NOTCH activation that could have gone un-noticed with longer-term approaches. Our finding does not exclude a canonical role for NOTCH. In fact we have shown that in somites, NICD also enters the nucleus where it presumably triggers transcription together with RBPJ^4, 5^. However, the phenotypical consequences of its nuclear functions in this tissue are unknown.

In conclusion, this study provides novel evidence for a transcription-independent function for NICD in somites and establishes AKT as a novel key player in the signaling cascade that leads to the formation of the early myotome in vertebrates

## MATERIAL AND METHODS

### Electroporation

The somite electroporation technique that was used throughout this study has been described elsewhere^2–4, 37^. Briefly, we targeted the expression of various constructs to the dorso-medial portion of newly formed interlimb somites of Hamburger–Hamilton (HH^38^) stage 15–16 chick embryos (24–28 somites). We have previously shown that this technique allows the specific expression of cDNA constructs in the DML, and that fluorescent reporters (e.g. GFP) are detected 3 hr after electroporation in this structure^39^. We have analyzed all embryos 6 hrs after electroporation, implying that the molecules under study here have been acting during a narrow timeframe (about 3 hrs).

### Expression constructs

#### The following constructs have been previously published

The CAGGS-H2B–RFP (provided by Dr. S. Tajbakhsh^4^) contains a fusion of Histone 2B with RFP, downstream of the CAGGS strong ubiquitous promoter (CMV/chick β-actin promoter/enhancer).

The CAGGS–EGFP^4^ contains the CAGGS promoter followed by the EGFP reporter gene.

The dominant negative form of SNAI1 consists of the chicken SNAI1, in which the repressor domain was replaced by the VP16 activator domain of the Herpes simplex virus^29^.

CAGGS HA-NICD-6xMyc contains the chicken NICD1 (aa 1763-2561) of the chicken NOTCH1, flanked with HA and 6xMyc downstream of the CAGGS promoter^4^.

The GSK3 biosensor (pCS2 GFP-GSK3-MAPK, Addgene #29689) contains a CMV promoter upstream of a GFP molecule followed by a polypeptide tail that contains 3 GSK-3β phosphorylation sites, a priming site for MAPK/Erk and a site for the binding of E3 polyubiquitin ligases^40^.

HES1–d2EGFP (provided by Dr. R. Kageyama) contains the HES1 mouse promoter followed by a destabilized d2EGFP^41^.

The membrane tagged NICD was constructed using Gibson assembly (NEB) with the signal peptide FGFR2 (from *Danio rerio*), the extracellular and transmembrane domain of human CD4 (both obtained from Dr. J Kaslin) followed by the HA tagged NICD previously described, cloned either into the bidirectional vector tetracycline responsive vector or into the pCX-Myc described above.

The H2B-BFP has been made by replacing the RFP from the CAGGS-H2B–RFP described above by a TagBFP (Evrogen) ^5^.

CAGGS-BFP has been made by cloning the TagBFP (Evrogen) into the pCAGGS vector.

#### The following plasmids were constructed

The four mouse NICD have been made by cloning them (addgene 20183, 20184, 20185 and 20186) using Gibson simple fragment assembly with pCAAGS HA-NICD-6xMYC (described in^4^) as a backbone, opened with ClaI and BglII restriction enzymes.

The deletion mutants of the cNICD1 and the RAM domain have been constructed according to the description domains^42^, by deletion of the pCAAGS HA-NICD described in^5^ with a Gibson double fragment assembly with the XbaI enzyme.

The delta CD4-NICD have been constructed as the delta NICD, using the pCAAGS CD4 HA-NICD-6xMYC^5^ as a backbone. The CD4 membrane-tethering tail contains the signal peptide FGFR2 (from *Danio rerio*), followed by the extracellular and transmembrane domain of human CD4 (both obtained from Dr. J. Kaslin).

The AKT variants have been constructed by Gibson simple fragment assembly with pCAAGS HA-NICD-6xMyc described in^4^ as a backbone, open with ClaI and EcoRI. The AKT variants were obtained from addgene 39531, 16243 and 17245.

The CD4-AKT variants have been constructed as the AKT variants, using the pCAAGS CD4 HA-NICD-6xMYC^5^ as a backbone.

### Immunohistochemistry, in situ hybridization

Whole-mount antibody staining was performed as described^3^. The following antibodies were used: rabbit polyclonals directed against chick myogenic regulatory factor MYF5 (obtained from Dr. B. Paterson^43^; 1/200); and anti-RFP (Abcam #62341, 1/1000); chicken polyclonal antibody against EGFP (Abcam #13970, 1/1000). Mouse monoclonal antibodies against c-myc-tag (Abcam #32072, 1/200), HA-tag (CST, #2367S, 1/100). In situ hybridization was performed as described^29^. Probes for AKT1, 2 and 3 were isolated by RT-PCR of total RNA extracted from chicken embryonic E19 brain and E14 muscle tissues mixed together. The primers designed for amplification were derived from GenBank chicken AKT1-3 sequences. Amplified fragments (about 600 bp each) were cloned into the pGEM-T vector and sequence-verified.

### Confocal analyses

Dorsal views of somites shown in all figures are projections of stacks of confocal images taken using a 4-channel Leica SP5 confocal microscope (Leica). Confocal stacks of images were visualized and analyzed with the Imaris software suite (Bitplane). Cell counting was performed using the cell counter plugin (Kurt De Vos, University of Sheffield) within the ImageJ software^44^.

### Quantifications and statistical analyses

Electroporation results in the transfection of a portion of the targeted cell population, which is variable from embryo to embryo. To precisely evaluate the phenotypes obtained after electroporation of cell-autonomously acting cDNA constructs, the number of positive cells was compared to the total number of electroporated cells, recognized by an internal fluorescent reporter construct. On average, more than 700 cells were counted per point and the corresponding quantifications are shown in all figures.

To determine the fluorescence intensity of electroporated cells, the surface of electroporated cells were rendered manually with Imaris software. The mean intensity for each cell and each channel in three-dimensions was collected for statistical analyses. Statistical analyses were performed using the GraphPad Prism software. Mann–Whitney non-parametric two-tail testing was applied to populations to determine the P values indicated in the figures. In each graph, columns correspond to the mean and error bars correspond to the standard deviation. ***p value < 0.001, extremely significant; **p value 0.001 to 0.01, very significant.

### Co-Immunoprecipitation

Two and a half day old non-electroporated (WT) chick embryos or embryos electroporated in the DML with HA-NICD-6xcMyc (NICD) or HA-CD4-NICD-6xcMyc (CD4-NICD) were incubated for 6hrs at 37°C. After screening, somites of approximately 40 embryos per condition containing electroporated (NICD and CD4-NICD) and non-electroporated (WT) DML cells were mechanically dissected under the microscope, homogenized using 26 gauge syringes and lyzed for 1h at 4°C using lysis buffer present in the coIP kit (Active Motif Co-IP kit). Cell Lysates were then centrifuged for 10min at 4°C at 14000 rpm and supernatants were dosed using Bio-Rad DC protein assay (DC protein assay kit, Bio-Rad, 500-0116). Cell lysates were subjected to co-IP experiment as described by Active Motif universal Magnetic coIP Kit (ref: 54002). Briefly, 1mg of total protein extracts were incubated with c-Myc antibody O/N at 4°C. The next day, 30uL of Magnetic beads were added to the mix and incubated for 1hr at 4°C. Negative control consists of NICD or CD4-NICD electroporated lysates with only magnetic beads addition. Afterward, beads were collected using a magnetic stick and supernatants were removed. The beads were washed for up to 5 times using the Co-IP/ wash Buffer in the Kit. Lastly, beads were resuspended with 2X Laemmli loading buffer and heated for 10min at 95°C before migration on a 10% SDS-page gel and transfer into a PVDF membrane (Millipore, IPVH00010) for western blot analysis.

### Western blot

For co-IP inputs lanes, 30 *µ*g of total protein extract samples and co-IP eluates resuspended with 2x Laemmli buffer were separated on a 10% SDS-PAGE gel, then transferred onto PVDF Immobilon-P membranes (Millipore, IPVH00010). After transfert, membranes were blocked with 5% of non-skimmed milk diluted in TBS-T (Tris-buffered saline (50 mM Tris, 150 mM NaCl, pH 7.4) + 0.1% Tween 20) and incubated overnight at 4°C with 1/1000 diluted primary antibodies including c-Myc (abcam; ab-32), GSK3β (cell signaling technology; cs-9832), AKT (cell signaling technology; cs-9272S) and GAPDH (cell signaling technology; cs-2118). Membranes were washed 3 times in TBS-T for 15 min and incubated for 1hour at RT with horseradish peroxidase-conjugated secondary antibodies (1/10 000). After 3 washes of 5 minutes each in TBS-T, membranes were incubated with enhanced chemiluminescence reagents (Amersham ECL Prime, GE Healthcare; Ref: RPN2232) and revealed by Chemiluminescence detecting machine from Thermo Electron Corporation (serial number: 392-562L (RS232C)).

#### SUPPLEMENTARY FIGURE LEGENDS

**Figure 2 - figure supplement 1:**
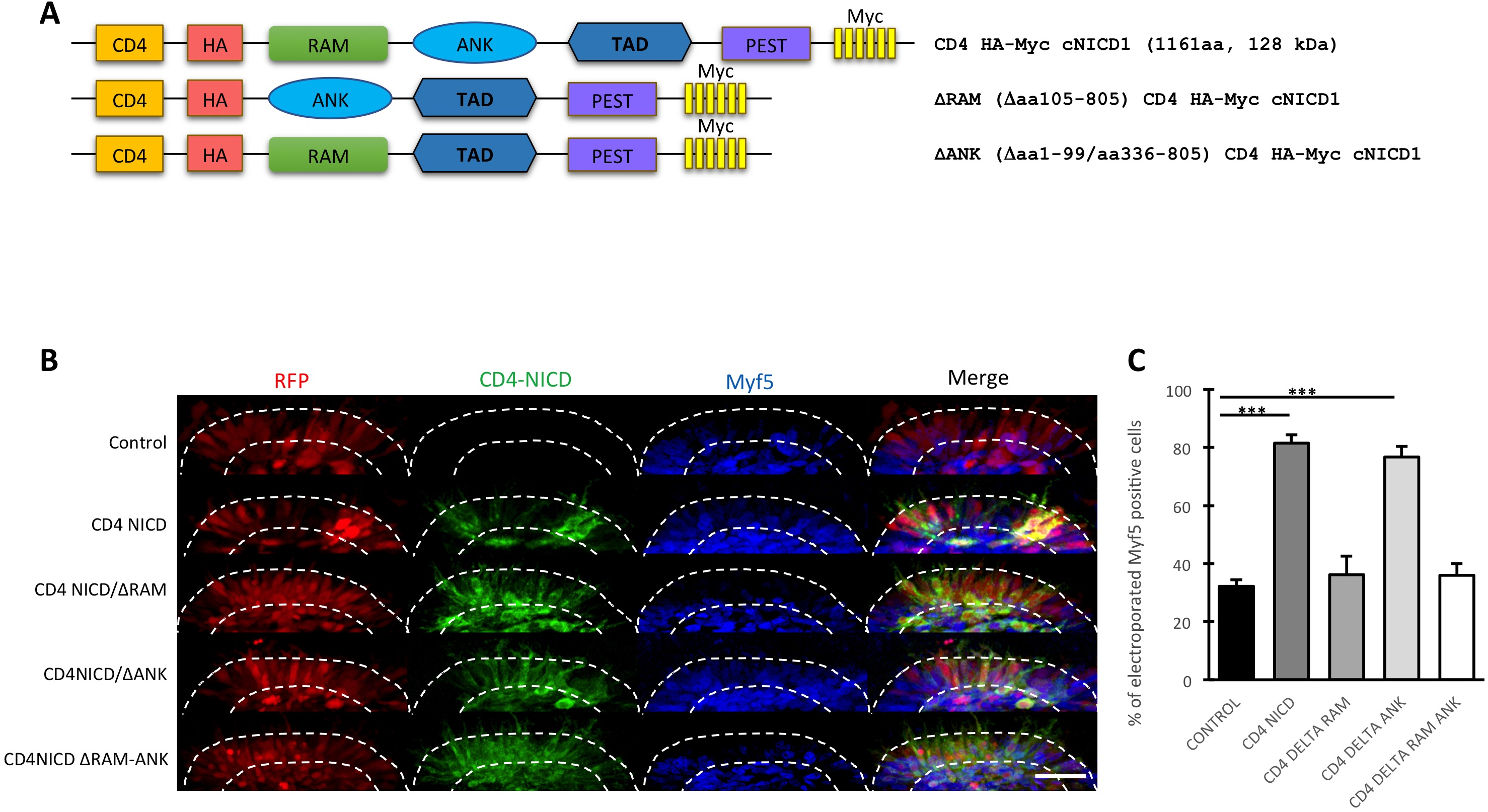
A membrane-tethered NICD depleted of its RAM domain is unable to activate Myf5 expression. (**A**) Schematics of the NICD mutants used in this experiments. All NICD constructs are tagged and attached to the membrane by fusion to the transmembrane domain of the human CD4. Indicates size of CD4 NICD1 (1161 aa) takes the tags and the CD4 membrane attachment into account. Amino acid numbering is derived from the chicken NOTCH 1 (GenBank accession # XP_015134811.1). **(B)** Confocal stacks of somites 6h after electroporation of mutant forms of a membrane-tethered (CD4)-NICD (in green), together with an electroporation marker (in blue). MYF 5 expression is in blue. **(C)** Bar charts showing that CD4-NICD and a CD4-NICD deleted of its ANK domain similarly activate MYF5 expression (81% and 77% of electroporated cells are MYF5+, respectively), compared to control embryos (32%). CD4-NICD deleted of its RAM domain or of the RAM and ANK domains are unable to activate MYF5 (36% in both conditions). ***:p<0,001. Scale bar: 50*µ*m

**Figure 3 - figure supplement 1:**
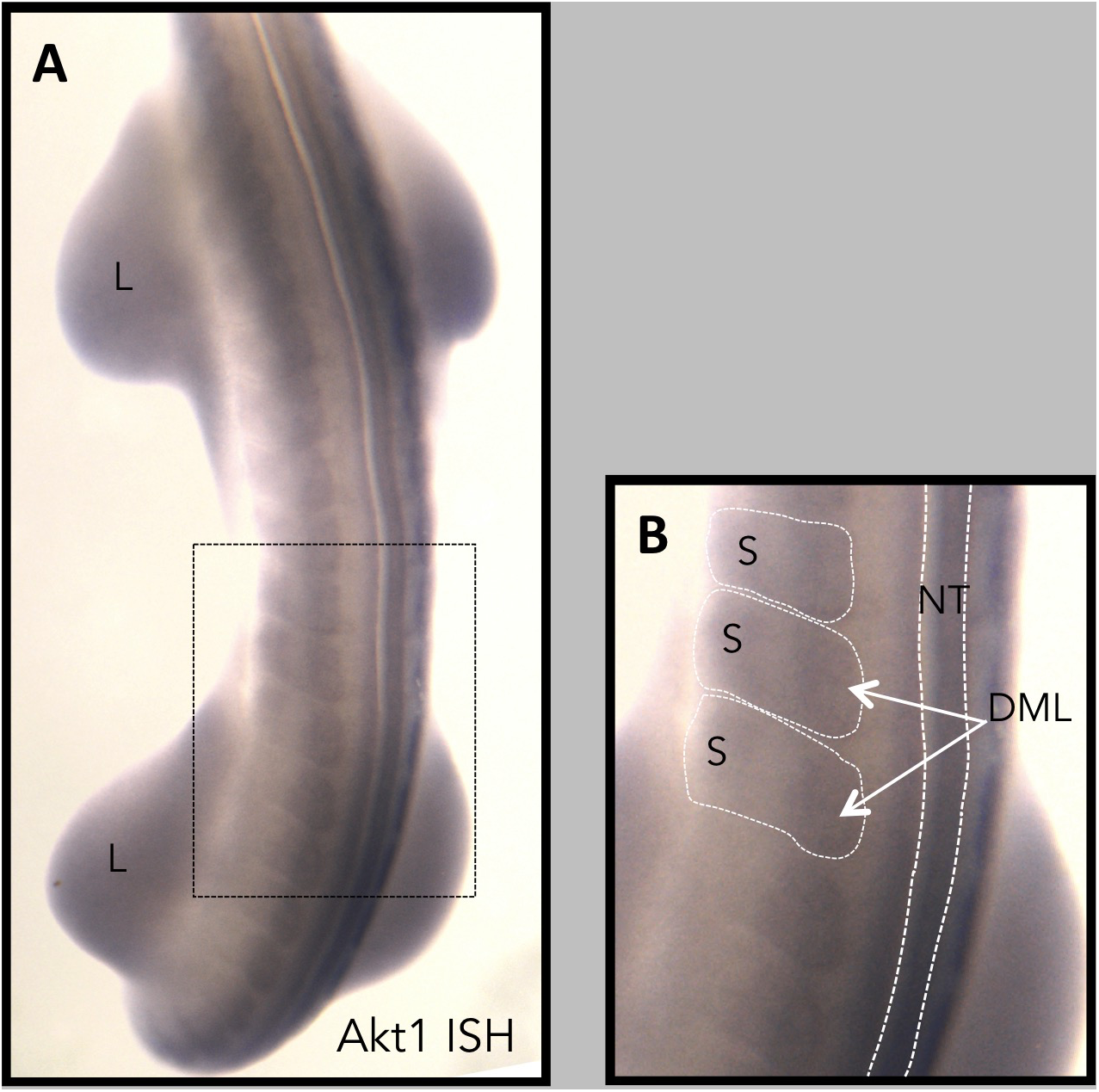
AKT1 is expressed in the DML of chicken somites. **(A,B)** Whole mount in situ hybridization on E 3.5 day old chicken embryos (Hamburger and Hamilton stage 21) with an AKT1-specific probe. Widespread expression is observed throughout the embryo, e.g. the limbs (L), the neural tube (NT) and the somites (S). The DML of all somites expresses Akt1 (arrows). **(B)** is an enlargement that corresponds to the boxed area in A.

## ACKNOWLEDGMENTS

We thank Monash Micro Imaging (MMI) and the Centre d’Imagerie Quantitative Lyon-Est (CIQLE) for imaging support; Profs. P. Currie and L. Schaeffer for critical reading of the manuscript. We thank Matthieu Marfoglia and Julie Melendez for their help with experiments. This work was funded by grants from the National Health and Medical Research Council (NHMRC, Australia), by the Australian Research Council (ARC, Australia) and by the Association Française contre les Myopathies (AFM, France). The Australian Regenerative Medicine Institute is supported by grants from the State Government of Victoria and the Australian Government.

The funders had no role in study design, data collection and interpretation, or the decision to submit the work for publication.

The authors declare no competing financial interest.

